# Linked-read analysis identifies mutations in single cell DNA sequencing data

**DOI:** 10.1101/211169

**Authors:** Craig L. Bohrson, Allison R. Barton, Michael A. Lodato, Rachel E. Rodin, Vinay Viswanadham, Doga Gulhan, Isidro Cortes, Maxwell A. Sherman, Lovelace J. Luquette, Minseok Kwon, Michael E. Coulter, Christopher A. Walsh, Peter J. Park

## Abstract

Whole-genome sequencing of DNA from single cells has the potential to reshape our understanding of the mutational heterogeneity in normal and disease tissues. A major difficulty, however, is distinguishing artifactual mutations that arise from DNA isolation and amplification from true mutations. Here, we describe <u>li</u>nked-read <u>a</u>nalysis (LiRA), a method that utilizes phasing of somatic single nucleotide variants with nearby germline variants to identify true mutations, thereby allowing accurate estimation of somatic mutation rates at the single cell level.

## Introduction

Comprehensive profiling of genetic mutations by whole-genome sequencing (WGS) has aided in answering fundamental questions in biology and medicine. If applied at the single-cell level, it has the potential to allow dissection of genetic heterogeneity at the highest resolution, allowing one to study mutational processes in greater detail than ever before. While single-cell RNA-seq can now be used to inexpensively assay thousands of cells to characterize small or rare cell populations, it cannot reliably be used to link expression heterogeneity with genetic heterogeneity. With the poor capture efficiency of published protocols (around 10% for many) and a high-level of technical noise, its sensitivity is too low for identifying mutations in most genes^1^; it also cannot profile non-coding regions. For DNA sequencing, the reduced cost and novel protocols for generating a single-cell library has enabled single-cell DNA sequencing (scDNA-seq) studies, including studies of exomes,^2–4^ low-coverage whole genomes for copy number variation (CNV) analysis,^5^ and more recently higher-coverage whole genomes.^6^ Widespread application of single-cell whole-genome sequencing (scWGS), however, has been hindered mostly by the errors associated with whole-genome amplification (WGA).

There are three main WGA protocols in use: degenerate oligonucleotide primed PCR (DOP-PCR), multiple annealing and looping based amplification cycles (MALBAC),^7^ and multiple displacement amplification (MDA).^8^ Although DOP-PCR and MALBAC generate a reproducible coverage profile and have been used for detection of large-scale copy number variation,^9,10^ their coverage is confined to a subset of the genome. For somatic single-cell SNV detection, MDA tends to produce the lowest false positive and negative rate, owing to the high fidelity of ϕ29 polymerase (1 error in 1e6-7)^11^ and a generally low allelic dropout rate and high genomic coverage. As such, MDA has become the most common WGA method in studies aiming to identify somatic SNVs (sSNVs).^9,10^

Nevertheless, MDA still suffers from issues that can confound studies aiming to call sSNVs. Despite its relative fidelity, ϕ29 is expected to produce hundreds to thousands of polymerase errors in the first replication of the genome alone. Additionally, the original DNA, if damaged during cell lysis or subsequent steps, may be unfaithfully copied early in amplification. In particular, heat is known to induce cytosine deamination,^12–14^ and deaminated cytosine created during cell lysis may introduce a substantial burden of artifactual C>T mutation calls. As an exponential amplification process, MDA will cause early errors to propagate. While theoretically these artifacts will have a lower variant allele fraction (VAF) than expected for an SNV, high variance in VAF due to allelic dropout and amplification bias may cause a fraction of artifactual calls to reach VAFs comparable to those of true mutations in the final amplification product.

This high level of technical noise has made the validation of putative SNVs important. Given the unavailability of the original, unamplified DNA sample after MDA, past studies have approached this issue by confirming that candidate SNVs from individual cells are present in other cells from the same organism. This can be done using an orthogonal technology such as ddPCR or amplicon sequencing in bulk DNA or looking for SNVs that were shared across multiple MDA-amplified single cells.^6,7^ While this strategy is viable for the validation of mosaic or subclonal SNVs, it cannot validate those detected in only one cell (singletons). This precludes analysis of post-mitotic SNVs and SNVs shared by arbitrarily small populations of cells.

## Results

Here, we present single-cell <u>li</u>nked-read <u>a</u>nalysis (LiRA), a method that aims to provide robust validation for a subset of candidate sSNVs occurring close to germline heterozygous SNPs (gHets) (Figure 1). The key insight underlying LiRA is that false positive sSNV calls (FPs) are derived from factors specific to one strand of DNA, while true positive sSNV calls (TPs), as fixed mutations, are derived from both strands of (almost certainly) one chromosome. A single read or two mate-pair reads covering the genomic positions of a putative sSNV and a nearby gHet can distinguish these two scenarios. For TPs, the subset of spanning reads supporting an sSNV call should support one of the two gHet alleles, consistent with the presence of a true SNV on one chromosomal copy. We call these reads ‘concordant’ reads (Figure 1a). In contrast, at FP sites, a mixture of support for the FP alternate and reference allele is observed on reads supporting the linked gHet allele, consistent with the acquisition of the FP during amplification or from only one strand of unamplified DNA. For FPs derived from DNA damage, these reads, which we call ‘discordant’ reads (Figure 1a), originate from the unaffected strand of the same chromosome, and for polymerase errors, from faithfully copied strands of the same chromosome (Figure 1a-b).

**Figure 1.**
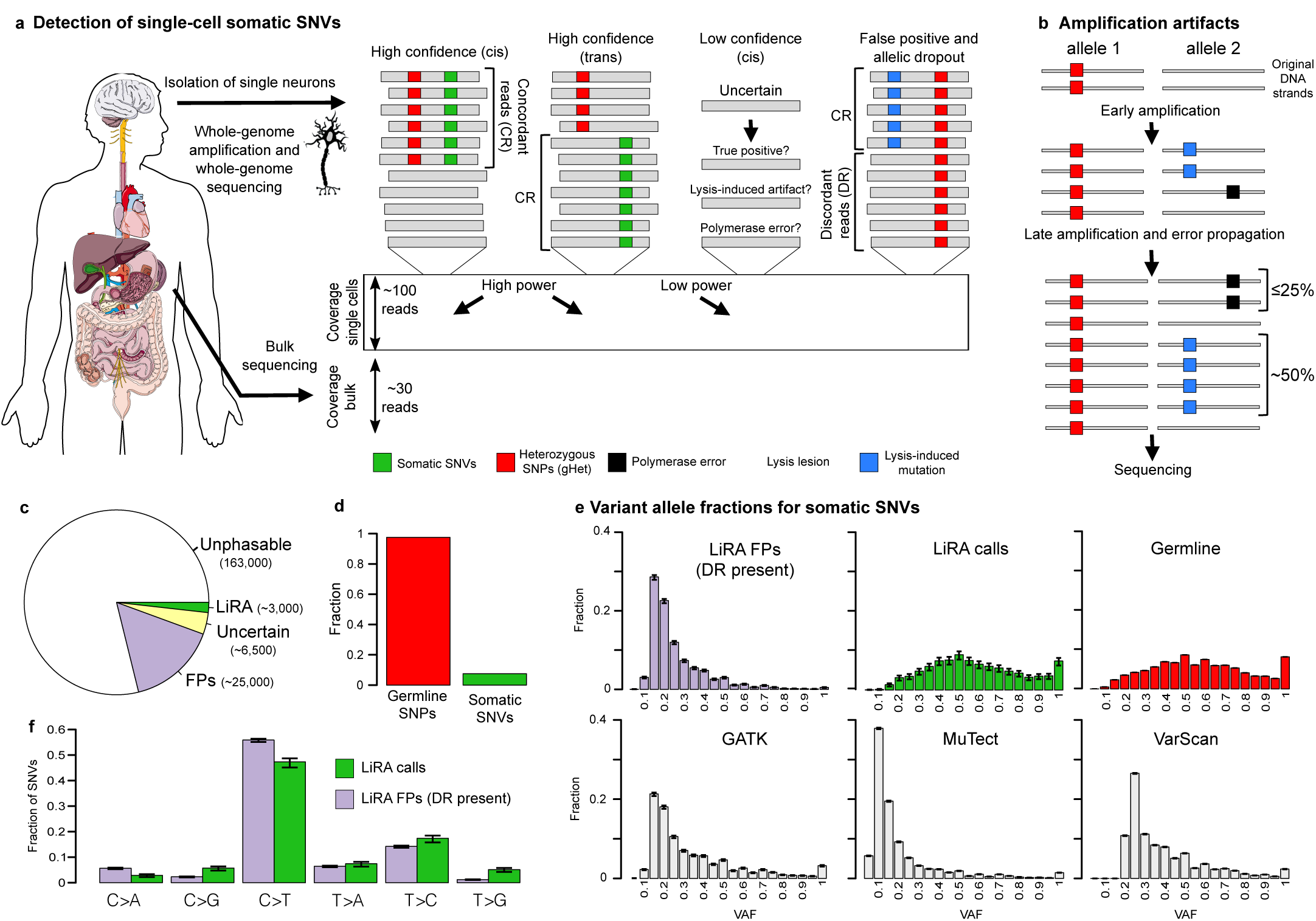
LiRA workflow, rationale, and variant properties. **(a) Overview of methodology for identifying false positive (FP) somatic SNVs (sSNVs).** ‘Concordant’ reads (CR) support the linked population SNP allele (gHet, red) marking the chromosome of origin for a somatic sSNV (green) and the somatic sSNV call. In *cis* linkage, SNV alternate alleles are always observed on the same set of reads; in *trans* linkage, the SNV alternate allele at one locus is always observed with the reference allele at the other. In contrast, ‘discordant’ reads (DR) for a lysis-lesion derived FP support a mixture of the spurious somatic call and reference. Due to the coverage non-uniformity introduced in amplification, power to distinguish these two scenarios is variable across the genome: when coverage is low, it is unclear whether a putative sSNV linked to a gHet with only concordant reads is derived from a true sSNV or an under-sampled FP. **(b) Model for how the linked read pattern specific to FPs arises from amplification of single-stranded DNA lesions or polymerase errors.** A lesion present on one strand of unamplified dsDNA is erroneously copied and creates an artifactual mutation early in amplification, but the unaffected strand is amplified as well, resulting in a mixture of support for the spurious mutation and reference on amplified DNA marked by the linked gHet allele. In an alternative scenario, φ29 polymerase mismatches a base while copying a strand of input DNA, resulting in the introduction of an artifactual mutation call linked with an adjacent gHet allele. As polymerase errors are introduced, after the first round of amplification at the earliest, they are expected to appear in ≤25% of gHet-linked reads, while lesions are expected to appear in ~50%. **(c) Classification of candidate sSNVs in LiRA.** Most sSNV candidates (est. ~163,000, 83%) cannot be analyzed using LiRA as they occur too far away from a gHet to be covered by the same read or mate pair. Of the fraction that can be analyzed (17%, ~34,500), the majority (72%, ~25,000) are identified as FPs due to the presence of at least one discordant read covering the sSNV position and each linked gHet. In the remaining subset, 6,500 (68%) of mutations are do not meet LiRA's quality thresholds due to a low coverage in bulk or single cell sequencing, and 3,000 (32%, 1.5% overall) are reported as LiRA sSNV calls. **(d) Differences in the phasing pattern observed between gHet-gHet and gHet-sSNV pairs.** 98% of linkable gHets are linked to a gHet with only concordant reads, whereas only 28% of linkable sSNV candidates are. Technical noise is expected to cause true heterozygous mutations, somatic or germline, to fail to link as expected, and analysis of gHets quantifies this effect as disqualifying only ~2% of true heterozygous mutations from analysis **(e) Comparison of the variant allele fractions (VAF) for different sSNV call sets** The VAF distributions are very similar for LiRA and germline calls, whereas those for standard genotyping methods are similar to the distribution of FPs identified by LiRA. 99% CI are shown. **(f) Comparison of mutation frequencies for LiRA calls and FPs as identified by LiRA**. C>A and C>T are enriched in FPs but by a small amount. 99% CI are shown.

In LiRA, we first utilize GATK^15^ or another variant caller with unrestrictive parameters to identify as many potentially somatic SNVs as possible, increasing our sensitivity. Candidate sSNVs are identified as any variants that are not present in matched bulk sample, and gHets as any population-polymorphic heterozygous calls made in bulk. We identify SNV-gHet pairs that are simultaneously supported by the same read or mate-pair, tracking pairs that have discordant reads. SNVs that are found exclusively in pairs with discordant reads are filtered as FPs.

We applied our method to single neuron data (sequenced at ~45X) from three phenotypically normal individuals.^6^ We find that a substantial portion (17%, Figure 1c) of SNV candidates can be subjected to this analysis and many identified as FPs (72% of the subset, Figure 1c-d). To assess specificity, we apply the same filter across gHet-gHet pairs and find that 98% of all gHets were retained. The high retention rate of gHets suggests that filtering sSNVs in this manner specifically removes FPs, and the relatively low retention of sSNVs suggests that the burden of FPs in the overall sSNV call set was high (at least 72%), indicating that the standard genotypers cannot be used in to call sSNVs without heavy filtering and validation.

After this filtering, LiRA thresholds the remaining set of sSNV calls based on a measure of quality called composite coverage (CC), defined as the minimum of spanning read depth in bulk and single-cell sequencing data (Figure S1; supplemental methods). As bulk coverage increases, it becomes more likely that an observed sSNV is not a missed germline event. As single-cell coverage increases, so does confidence that discordant reads are truly absent in the MDA amplification product, and not simply under-sampled. When single-cell coverage is low, FPs may appear linked without discordant reads to nearby gHets because of insufficient sampling, but as coverage increases this scenario becomes less and less likely. LiRA approaches this issue by finding a coverage threshold that controls the collective FP rate at a tolerable level (here 10%). Of the remaining SNV candidates, those with subthreshold support are called as uncertain, while those with support equal to or exceeding the threshold are called as TPs (Figure 1c-d).

To determine an appropriate threshold, LiRA takes advantage of the fact that in the absence of FPs, the estimated genome-wide somatic mutation rate should not depend on the quality of SNV calls. LiRA models the relationship between the somatic mutation rate and quality as the mixture of an exponentially decaying error component and an approximately constant true mutation component (supplemental methods). We find that this model fits the data well with a decay rate of 1/2 (Figure S2), which is consistent with DNA lesion-derived FPs comprising most of the error signal after filtering non-concordant candidate sSNVs. The utility of the model lies in the fact that the fitted true mutation component gives an estimate of the genome-wide somatic mutation rate, and its value relative to the error component gives an estimate of the FP rate at each level of SNV quality. This information, combined with the observed distribution of quality values, can be used to assign FP probabilities to individual mutations, and to ascertain the aggregate FP rate expected across a set of calls when thresholding at different quality values. Importantly, this model is fit for each cell individually, and thus can account for variable artifactual burdens, allowing control of the false positive rate across all cells at a specified level while maximizing sensitivity. In the Lodato *et. al.* (2015) data, this proved to be an important consideration, as the estimated artifactual burden among SNVs with low support (CC = 2), varied by 3-fold across cells (Table S1). Fitting the model for each cell also revealed that this measure of quality was uncorrelated with the estimated total number of somatic mutations per cell, consistent with LiRA correctly identifying and separating the errors and true mutations (Table S1). Finally, we found that our estimated genome-wide somatic mutation rates in prefrontal cortical neurons were generally consistent with those found by a previous study^16^ that measured the rate of mutation in the frontal cortex and avoided use of single-cell sequencing or WGA (Table S1). This suggests that our method accurately accounts for the heterogeneity in power introduced through MDA-related coverage non-uniformity and LiRA's specific analysis of regions around gHets.

Although only a tiny fraction (Figure 1c-d) of initial SNV calls are eventually called as TPs, we find LiRA retains surprisingly high sensitivity. Using genome-wide mutation rate estimates and the number of passing mutations for each cell, we estimate LiRA's sensitivity to be, on average, 9% on the Lodato dataset, detecting on average 83 somatic mutations out of an average extrapolated genome-wide rate of 946 somatic mutations per cell (Table S1).

To validate that LiRA was selecting the TP subset of candidate SNVs, we compared the variant allele fraction (VAF) distribution of LiRA sSNV calls to that of germline SNVs, as well as to the VAF distribution produced by GATK,^15^ MuTect,^17^ and Varscan^18^ for somatic calls (Figure 1e). Calls reported by LiRA produced a VAF distribution nearly identical to the germline distribution, whereas calls reported by other variant callers produced VAF distributions highly skewed towards low VAF calls.

We also compared the genomic context in which LiRA calls and FP mutations occurred, reasoning that if the LiRA calls were truly derived from double-stranded fixed mutations, the genomic context would be measurably distinct. The LiRA calls and FP mutations had significantly different abundances of each SNV type, consistent with the two sets originating from different underlying processes. C>G, T>A, T>C, and T>G mutations were all enriched in TPs, while C>T and C>A mutations were depleted (Figure 1f, S3). The largest enrichment was observed for T>G (4 fold) and the largest depletion for C>A (2 fold). To get a more detailed view, we compared the trinucleotide context for TP and FP mutation calls, which revealed larger differences in C>T and T>C mutations between the TP and FP sets (Figure 2). GCN>T and CTN>C mutations were depleted (1.9-3.9 fold) among true SNVs, while ATN>C mutations were enriched (1.9-9.8 fold). We also found that LiRA TPs are enriched relative to FPs at H3K36me3 marks in mid-frontal cortical neurons (Figure S4). Interestingly, this coheres with results suggesting that these regions are targeted by mismatch repair (MMR)^19^ and that in non-dividing cells MMR is mutagenic.^20^

**Figure 2.**
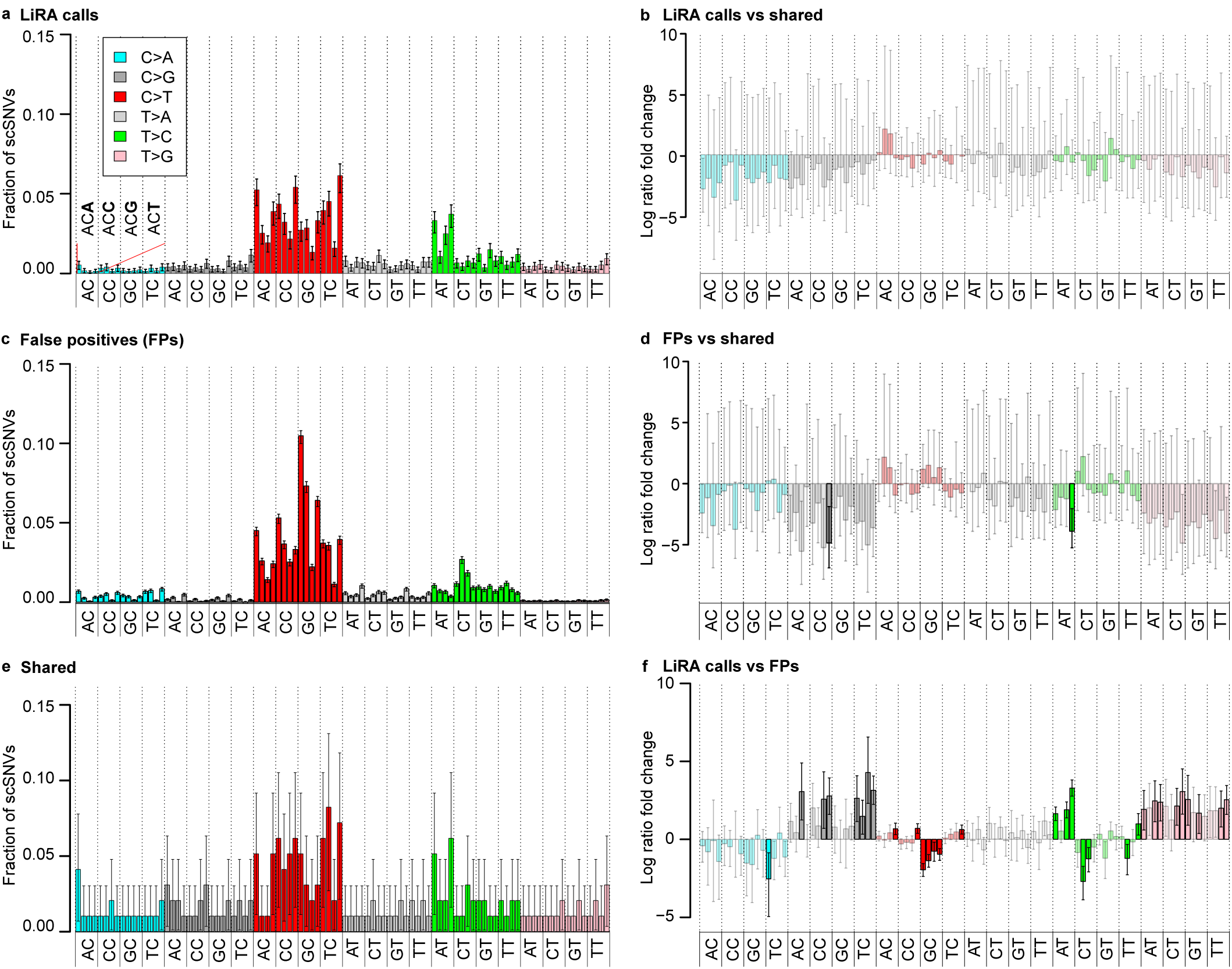
Comparison of the trinucleotide contexts of LiRA calls, FPs, and shared mutations. LiRA calls found with support in more than one cell (99% CI). **(a)** The trinucleotide contexts of LiRA TPs (99% CI). **(b)** Log-scale fold change in mutational abundance between LiRA calls and LiRA TPs (99% CI). No fold-changes reach significance. **(c)** The trinucleotide context of filtered FPs (99% CI). **(d)** Log-scale fold change in mutational abundance between FPs and shared mutations (99% CI). Significant changes are highlighted. **(e)** The trinucleotide contexts of shared (more than one cell) mutations (99% CI) **(f)** Log-scale fold change in mutational abundance between LiRA calls and FPs (99% CI). Significant changes are highlighted.

As further validation, we identified LiRA calls shared across cells and compared the trinucleotide context of this set to that of the non-shared LiRA call and FPs. A shared mutation was called if it was called by LiRA in one cell and had supporting reads in another cell. The abundance of shared LiRA calls in the Lodato et. al. 2015 data was low (7/2980, Table S2), and so we also looked for LiRA calls from Lodato et. al. 2015 in new neurons sequenced from UMB1465 (Lodato et. al. 2017, in submission). A surprisingly high number of mutations were shared between exactly one original neuron and one new (38 between cells 8 and p2b11; 50 between cells 43 and p2f06, Table S2), and this raised the number of shared mutations detected to 96/2980. With this increased number of observations, we found that there were no significant differences between the trinucleotide context for the private and shared LiRA calls (Figure 2b), but that the differences between LiRA FPs and shared calls mirrored those between the LiRA calls and FPs (Figure 2d and 2f). When we collapsed the trinucleotide context into a single nucleotide view to gain more power (Figure S3), the difference between LiRA calls/shared mutation and FPs became more obvious. The similarity in nucleotide context between private and shared LiRA calls and their mutual difference from FP calls suggests that LiRA accurately identifies mutations in single cells with high specificity.

## Discussion

While we find significant evidence consistent with LiRA calling a set of fixed, double stranded sSNVs, there are in theory error modes in amplification that would cause FPs to escape LiRA's filtering steps. LiRA relies on both strands of a single chromosome being subject to relatively even amplification. If present, strand dropout or severe nonuniformity in strand-specific amplification could cause DNA lesions or polymerase errors to appear as fixed mutations in single-cell sequencing data. While we cannot technically rule this out, the quality of the two-component model fit across cells (Figure S2) as well as other properties of LiRA calls presented suggest this process is of negligible effect size. In the error model, artifactual mutations are expected to halve in abundance with every increase of 1 in composite coverage, which suggests the errors that persist at the highest composite-coverage values are supported in the MDA amplification product with the same abundance of concordant and discordant reads. This is consistent with the scenario in which the strand hosting a DNA lesion and its complement are subject to even amplification. As noise is introduced, the error model is expected to fit poorly, as a large number of artifacts would stochastically acquire a much higher abundance of concordant reads. This would result in a much slower exponential fall in mutation rate with increasing composite coverage as well as a poor fit of the exponential error model with its fixed rate of decay. Additionally, we expect that the strand-displacement activity of ϕ29 should encourage even amplification of both strands or neither, as amplification of one strand simultaneously inhibits new degenerate hexamer binding to the strand that gets copied and catalyzes hexamer binding to the opposite strand by displacing it as single strand DNA.

Our results show that LiRA comprises an advance in SNV calling in single cells, especially with respect to mutations private to one cell. As we showed (Figure 1e), a standard variant caller with default parameters does not perform well on single-cell data, with the majority of the calls being false positives. Existing sSNV calling methods aim to identify shared or clonal mutations, often in the context of cancer, and provide limited support for analysis of mutations found in only one cell. Monovar^21^ marks sSNVs shared across samples but has very lenient criteria for such cases, and, while it provides private calls, lacks a way of assessing their quality. Single Cell Genotyper^22^ is developed specifically to cluster variants in single cells to produce clades and identify groups of cells with a shared genotype rather than produce individual calls. While both may be useful to those studying clones or tumor dynamics, they cannot aid in the analysis of private or very rare somatic mutations.

Although LiRA is designed to utilize only the linked subset of putative SNVs, we show that it is sensitive enough to accurately estimate the genome-wide mutation rate as well as to characterize mutational spectra and their properties. This new approach to single-cell analysis can provide a window into the mutational processes within a cell that might lead to new insights into cell aging, lineage, disease, and more.

## Acknowledgements

This work was mainly supported by NHGRI (T32HG002295) and NIMH (U01MH106883). Some figures used images from the Servier Medical Art PowerPoint Image Bank.

## Supplemental Figure Legends

**Figure S1.**
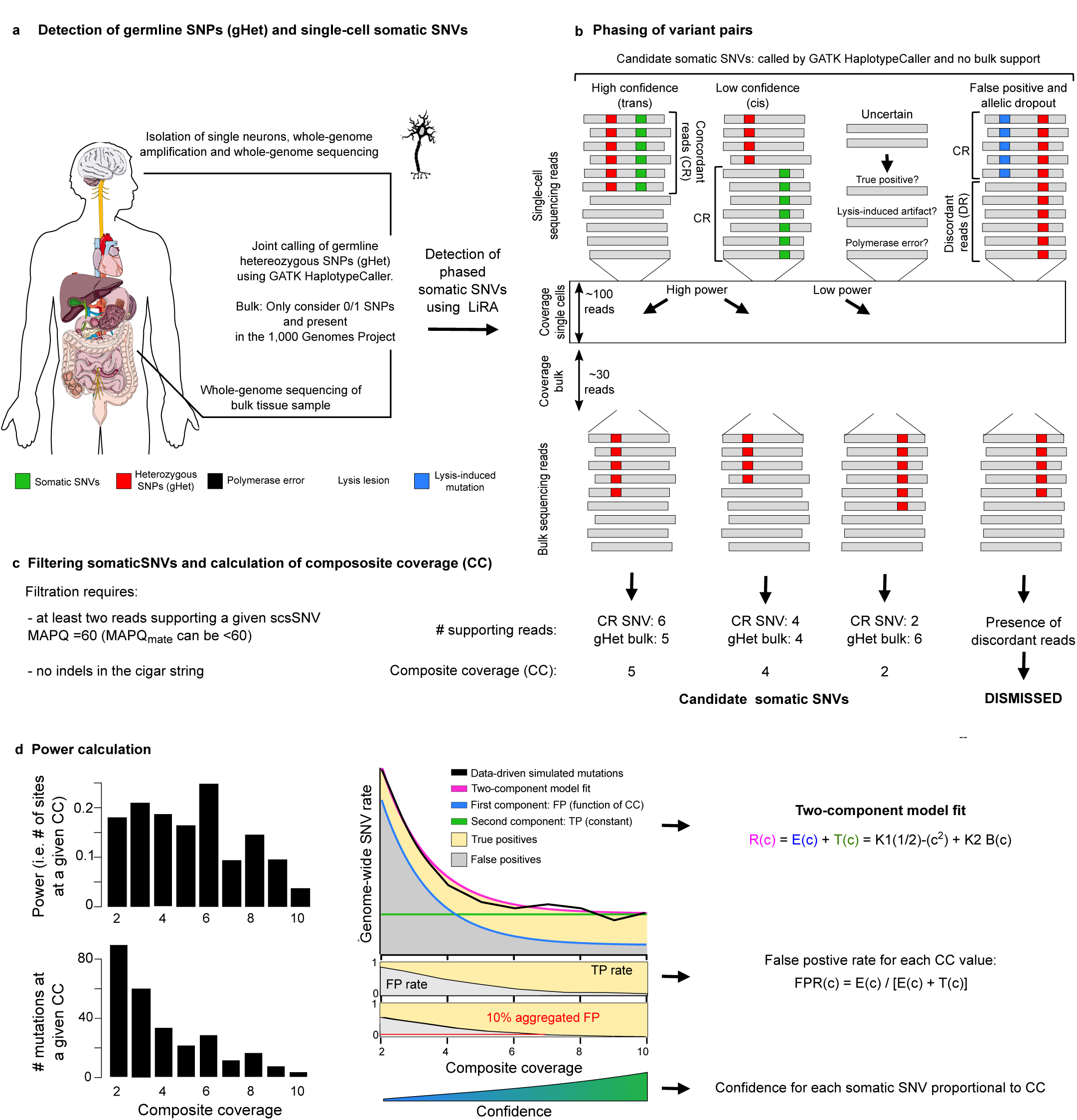
Detailed LiRA Workflow. **(a) Detection of germline SNPs (gHet) and single-cell somatic SNVs** LiRA uses germline heterozygous SNPs (gHets) found in bulk sequencing to evaluate nearby somatic SNV (sSNV) candidates. gHets are SNVs called with a ‘0/1’ genotype and annotated with a population frequency in the 1000 genomes database. **(b) Phasing of variant pairs** Candidate sSNVs are phased with nearby gHets. Power to distinguish true sSNVs from false positives scales with the minimum of single-cell and bulk sequencing depth only considering reads supporting the linked gHet allele. **(c) Filtering somaticSNVs and calculation of composite coverage (CC)** CC is measured for each sSNV candidate linked with only concordant reads to a nearby gHet as the minimum number of reads spanning the two loci in bulk and single-cell sequencing. Reads must have MAPQ=60 and no indel CIGAR operations to be considered. sSNVs linked only with a mixture of discordant and concordant reads are dismissed as likely FPs. **(d) Power calculation** Based on the frequency of sites at which an sSNV could have been detected with each CC value (power) and the frequency of CC values among candidate sSNVs, LiRA measures the somatic mutation rate as a function of CC. To the observed data (black; simulated), LiRA fits a two-component mixture model, wherein mutations either originate from errors (blue; E(c)) or real sSNVs (green; T(c)). E(c) decays exponentially as it becomes increasingly unlikely that an artifact will only be observed with concordant reads as CC rises. T(c) is derived from sampling the distribution of CC values for linked gHets and is approximately constant, as would be expected for true heterozygous variants. The relative magnitudes of T(c) and E(c) give estimates of the false positive rate (FPR) among variants with each value of CC. LiRA uses this information, along with the actual distribution of CC values over the candidate sSNV set, to choose a CC threshold that controls the overall false positive rate under a specified level (here 10%).

**Figure S2.**
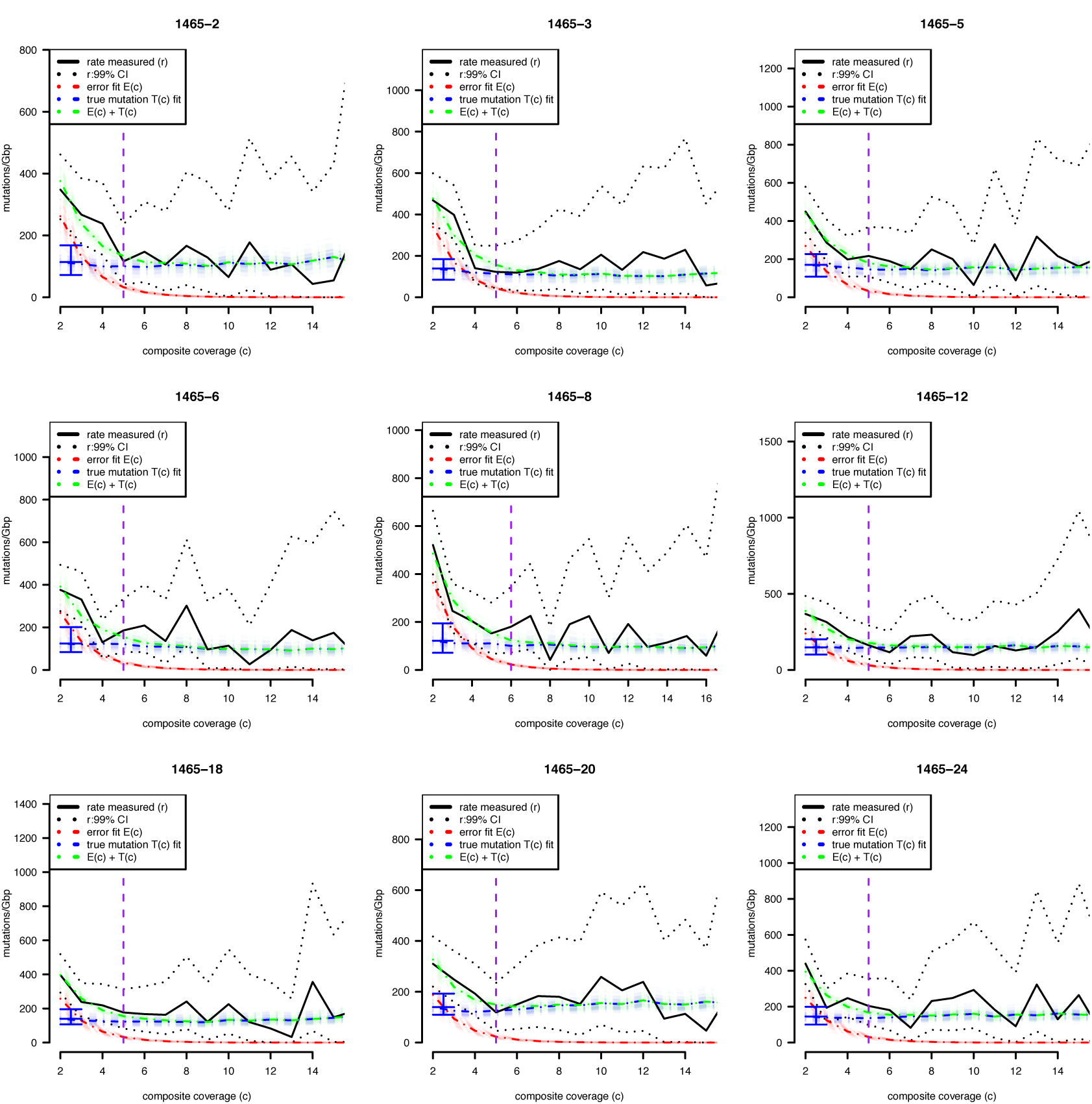
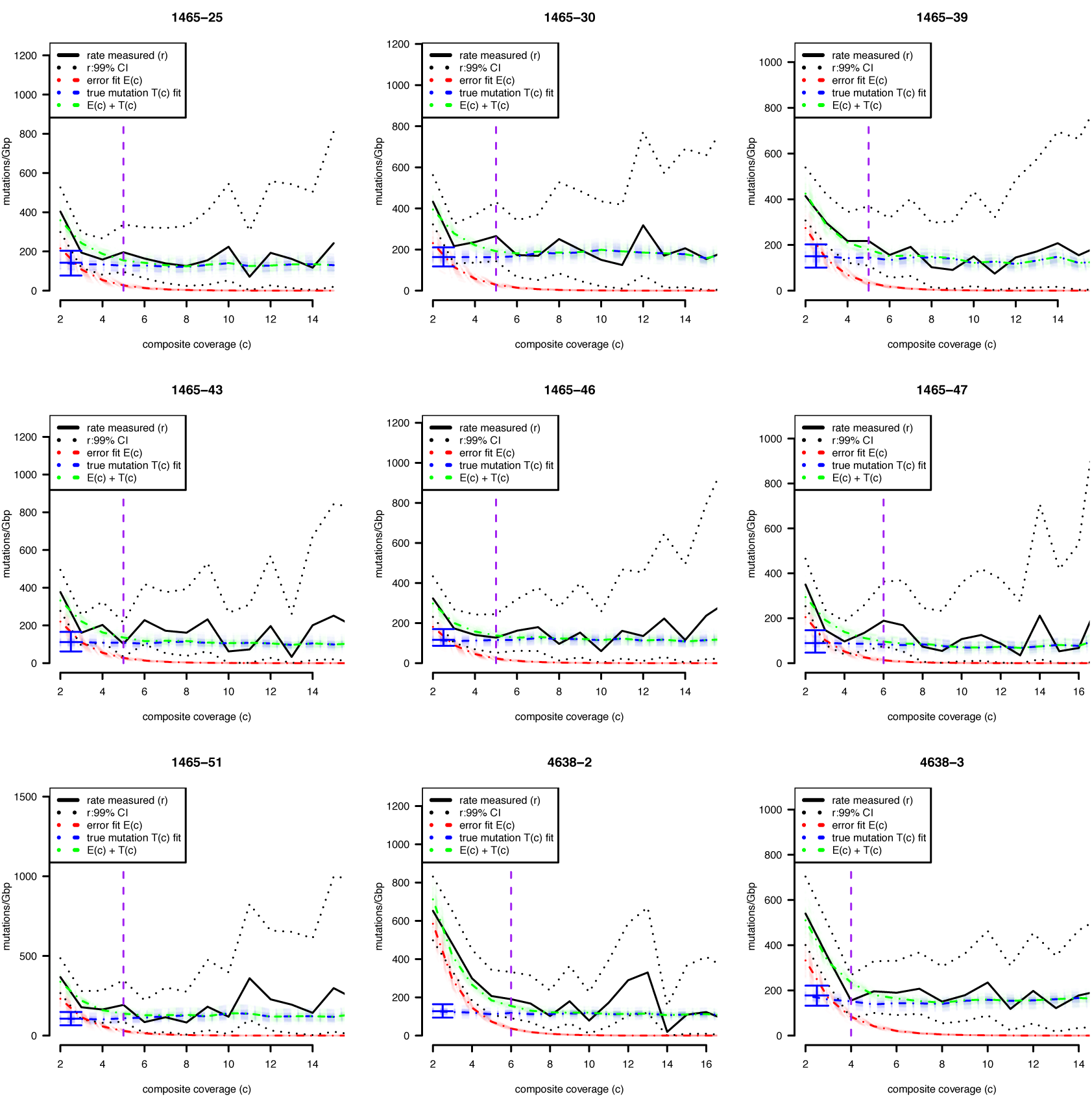
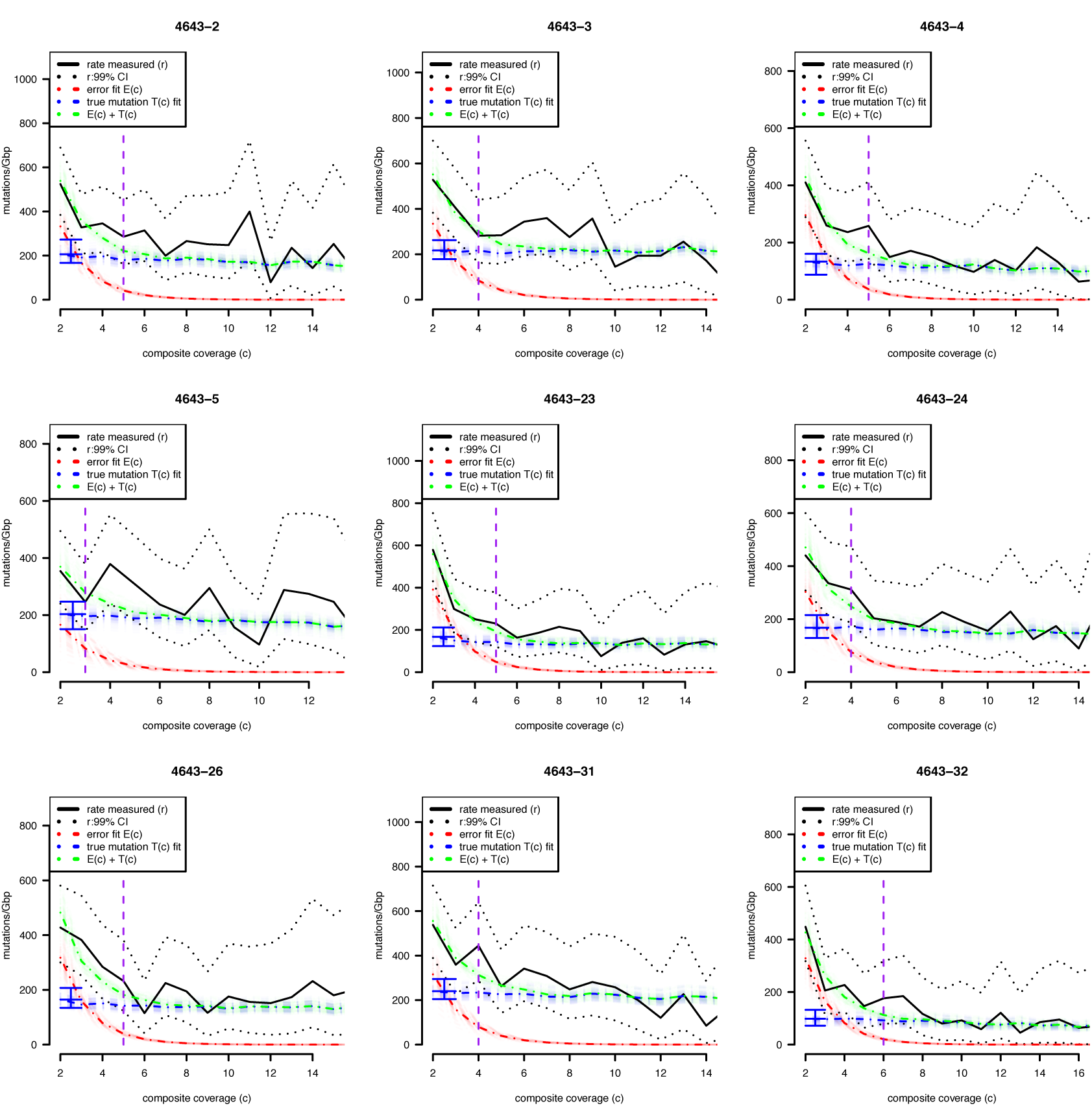
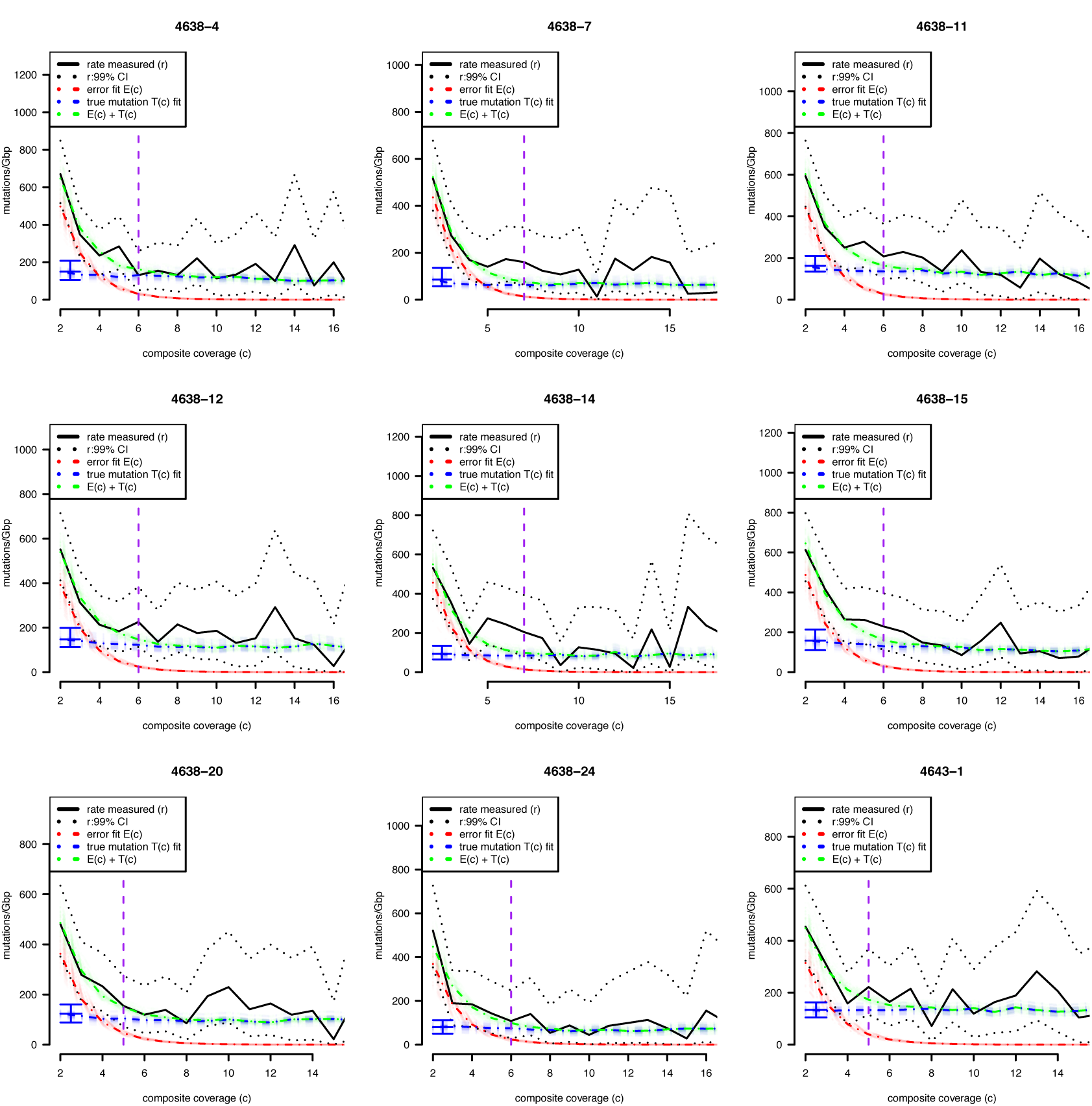
Two-component model fits for all neurons. Faded dotted lines show 100 fits obtained from simulating random draws from the beta distribution provided by the observed data (see methods), and the blue error bar shows a 98% confidence interval for sSNV rate.

**Figure S3.**
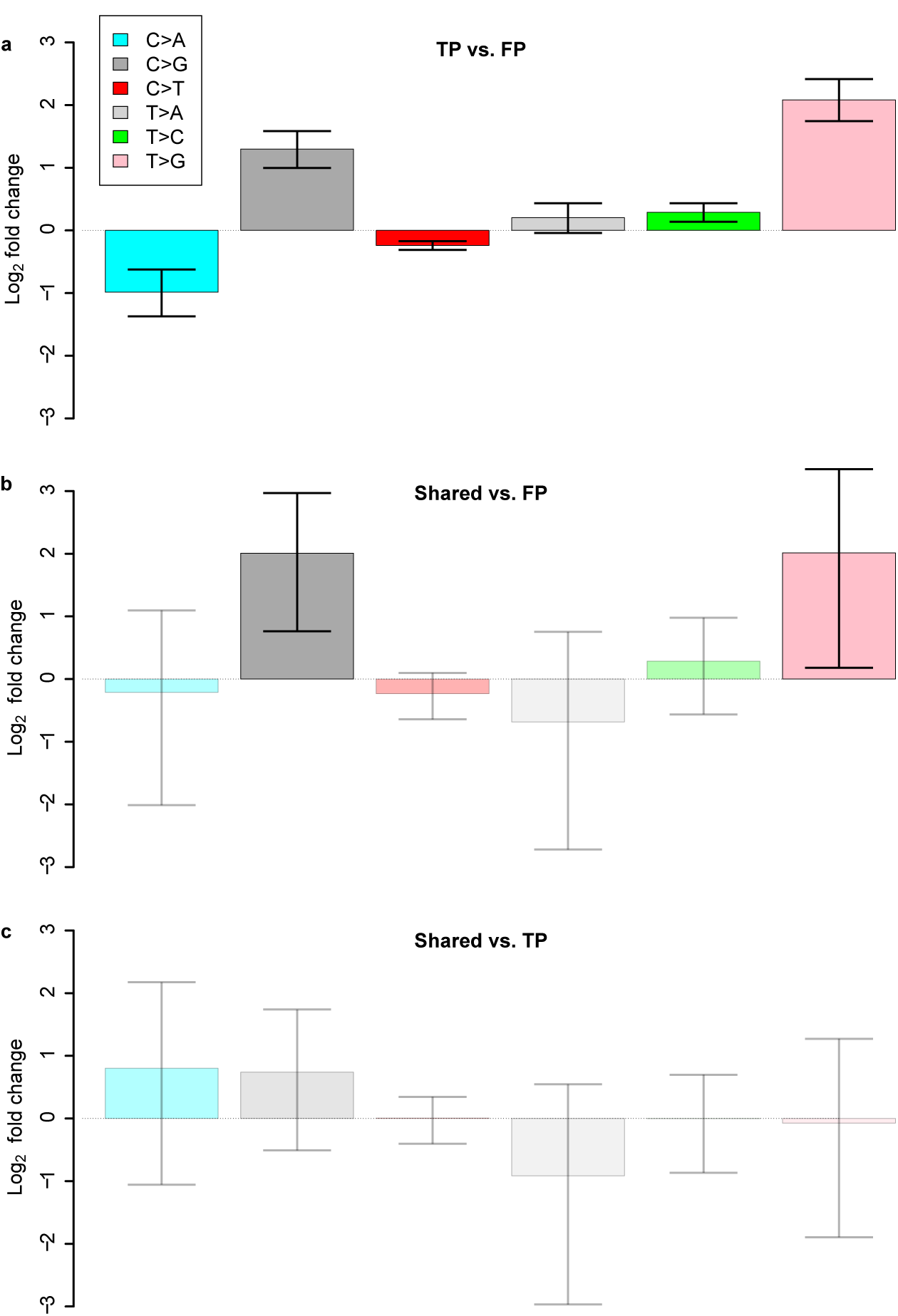
Comparison of SNV frequencies for LiRA calls, FPs, and shared mutations. LiRA calls found with support in more than one cell (99% CI) **(a)** Log-scale fold change in mutational abundance between LiRA singleton calls and LiRA FPs. **(b)** Log-scale fold change in mutational abundance between LiRA shared calls and LiRA FPs. **(c)** Log-scale fold change in mutational abundance between LiRA shared calls and LiRA singleton calls (99% CI).

**Figure S3.**
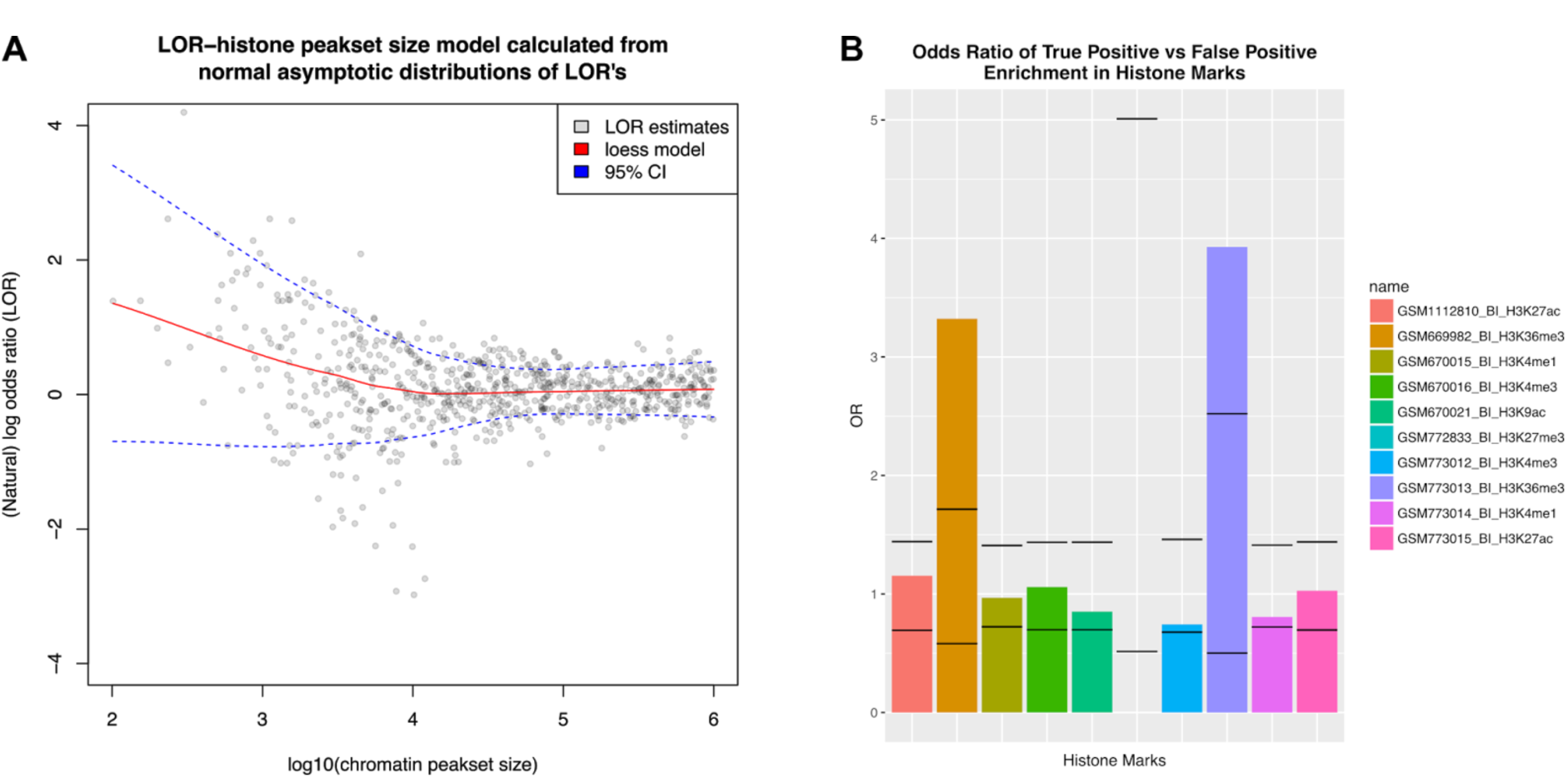
True positives called by LiRA are enriched in H3K36me3 in mid-frontal cortical neurons. **(a)** Null model showing the change in expected natural log odds ratio (LOR) of TP vs FP enrichment as the abundance of genomic intervals associated with random chromatin marks increases. 95% CIs are shown for the LOR. **(b)** Observed LOR's of TP's vs FP's in specific chromatin marks representing accessible chromatin or specific histone modifications in adult mid-frontal cortical neurons. LOR's for each mark are shown along with 95% CIs for a control peakset of comparable size as calculted in (a). TP's are enriched in genomic intervals with H3K36me3 modifications. Observed odds ratio for H3K27me3 could not be determined, as instances of true or false positives were not found in H3K27me3 owing to the histone mark's small peakset size (~10^3^).

## Supplemental Methods

### Variant calling of candidate single-cell somatic single nucleotide variants (scsSNVs) and population-polymorphic germline heterozygous sites (gHets)

The GATK Haplotype Caller best practices pipeline^1^ with default parameters was used to call variants jointly on single cell and bulk sequencing data. This yielded a large set of candidate sSNVs and gHets. Candidate sSNVs were identified as calls with no supporting reads in bulk and at least one supporting read in a single cell. High-confidence germline heterozygous sites were identified as variants found with population frequencies in the 1000 genomes database^2^ as annotated in the dbsnp147 database and called with a ‘0/1’ heterozygous genotype in bulk.

### Identification of candidate variants for LiRA analysis

scsSNV-gHet pairs and gHet-gHet pairs that had at least two reads or mate pairs supporting each variant locus were subject to analysis by LiRA. Included reads were required to have max mapping quality (60), to align concordantly (SAM flag 2), and to have no indel cigar operations.

### Phasing of variant pairs

Phasing was done by simple majority counting of reads from single cell (sSNV-gHet) or single cell and bulk (gHet-gHet) sequencing data. Paired variants *v* and *q* were *cis* linked with respect to *v* if spanning reads supporting *v* alt calls more frequently supported alt than ref at *q*. Otherwise, they were *trans* linked with respect to *v*. Importantly, this procedure defined the form of discordant and concordant reads for individual variants within variant pairs: discordant reads for *v* (*v*-discordant reads) were reads supporting the *v*-phased allele at *q* (alt for *cis*, ref for *trans*) but not the alt call at *v*.

### Computation of composite coverage and identification of discordant variants

For each variant pair **p** with variants *v* and *q*, the bulk phased coverage and single cell phased coverage for *v* (C_bulk:**p,***v*_, C_single-cell:**p**,*v*_) were measured as the number of spanning read pairs supporting the *v-*phased allele of *q* in bulk and single cell data, respectively. The composite coverage was computed as the minimum of these two values:

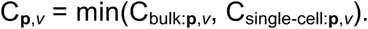

To obtain an overall composite coverage value for *v*, we took the maximum composite coverage value measured over all pairs of which *v* was a member:

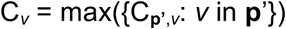

*v* was defined as a discordant variant if all pairs of which *v* was a member had at least one *v*-discordant read. *v* was defined as a concordant variant if it was not discordant. Discordant sSNVs were filtered as FPs, for figure 1d, discordant gHets comprise the 2% minority not found in a gHet-gHet linked pair with only concordant reads.

Over scsSNV-gHet pairs, the bulk phased coverage provided a measure of confidence that an scsSNV was not missed in bulk due to low power. The single cell phased coverage provided a measure of confidence over concordant variants that an scsSNV did not spuriously appear concordant due to under-sampling.

### Calculation of power by genomic position

First, all gHets were phased using SHAPEIT2.^3^ This yielded the chromosomal copy (1/2) of origin for the alternate allele of each gHet. Next, for each chromosome (*a*) of each gHet (*g*), the set of read pairs from bulk (R_bulk:*a*,*g*_) and single cell (R_single-cell:*a*,*g*_) covering *g* and supporting *a* were extracted. At each position *x* covered by reads in R_bulk:*a*,*g*_, the bulk phased power was measured as the depth of coverage in R_bulk:*a*,*g*_ at *x*. This was the bulk phased power for position *x*, using allele *a* of gHet *g* (P_bulk:*x*,*a*,*g*_). The single cell phased power (P_single-cell:*x*,*a*,*g*_) was computed similarly using R_single-cell:*a*,*g*_. The composite power for position *x* on chromosomal copy *a* with respect to gHet *g* was computed as the minimum of these two values:

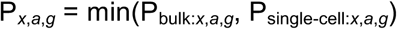

This procedure mirrored the composite coverage computation for non-variant sites. If there had been an sSNV *s* at *x* on chromosomal copy *a*, we would have computed the composite coverage for *s* in pair *s*-*g* as P_*x*,*a*,*g*_.

The overall composite power for position *x* on chromosomal copy *a* was computed as the maximum observed with respect to any gHet:

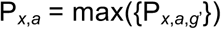

### Aggregate power calculation

Tabulating the output from the calculation of power by genomic position yielded counts (**A**) of the number of positions genome-wide with each possible value of composite power. These counts were adjusted from their raw values to account for two factors: loss of power due to non-artifact driven discordant read observations and loss of power due to the random occurrence of bulk-alternate reads supporting sSNV calls. As composite coverage increases, so does the likelihood that a discordant read will be observed due to technical noise. Similarly, the likelihood that a read from bulk sequencing data will support an sSNV increases. Both these events will reduce power to detect sSNVs by some fraction yet unaccounted for, since a single discordant read would, in principle, result in total loss of power at a given position. Our approach to this issue was to adjust each entry of **A** by a composite power dependent fraction (**f**_sd_) to account for stochastic discordance, and a fixed fraction for stochastic bulk support (*f*_bs_).

To compute *f*_sd_, we calculated the fraction of gHets found to be concordant as a function of composite coverage. As expected, this was generally high (fig 1e) but decreased as composite coverage increased.

To compute *f*_bs_, we calculated the fraction of gHets found to have no reads supporting a third allele, and set *f*_bs_ as this rate divided by two. While we expected there to be a dependence between the composite coverage and the rate of third allele observations, we did not see one, and thus used a fixed fraction instead of a composite-coverage dependent quantity.

We then adjusted **A**:

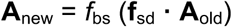

### Rate calculation and two-component model

To obtain estimates of and bounds on the mutation rate (**R**) at different composite coverage values we used a beta distribution with a uniform prior:

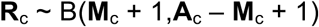

Where **M**_c_ is the number of mutations with composite coverage *c*.

Consistent with the idea that concordant candidate sSNVs with low composite coverage values are under sampled discordant variants, the mutation rate consistently (across cells) and rapidly decreased from its initial value at *c*=2 and approached a constant.

Rather than an arbitrary threshold for composite coverage by eye, we developed a method of estimating the false positive rate at different composite coverage values. Further, we used this to estimate the false positive rate across individual cells’ entire dataset when thresholding at a particular composite coverage value *c*\*. This allowed us to set a false positive rate constraint (<10%) and enforce it across multiple cells.

For each cell, we modelled **M** as the mixture of an error component (E) and a true component (T). The error component we fit had the form:

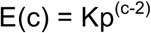

Visually, a decaying exponential appeared to fit the data well at low values, but this is also how we expected undersampled discordant variants to behave theoretically. If we assume an initial burden of K errors per gigabase at *c*=2, and that the probability of sampling a concordant read given a variant is truly discordant is *p*, then the error abundance as a function of composite coverage takes exactly this form.

We found that p = ½ resulted in good fits, and this suggested that the artefacts causing an excess of mutations at low composite coverage values originated from lesions present on the original DNA prior to any amplification. These were likely induced during cell lysis.

The “true” component T(c) is practically constant, but to improve the quality of fitting was actually computed using a bootstrapped set of germline variants. The procedure is as follows:

1. Randomly select a set of germline variants of size equal to the size of the scsSNV set (*c* > 2) from those found in gHet-gHet pairs, constraining the distance and cis/trans distribution to be as close as possible to that observed in the somatic.
2. Compute the rate using **A** and the composite coverage distribution over these gHet-gHet pairs.
3. Compute the bootstrap rate B(c) (the “true” component) by averaging over 100 instances.

Overall, the model R(c) = E(c) + T(c) = K_1_(1/2)^−(c-2)^ + K_2_ B(c) was fit using the R function nlm.fit, constraining K_1_ and K_2_ to be positive by imposing a large penalty on the objective function for K_1_ or K_2_ less than 0.

### Computation of false positive rate (FPR) and choosing a threshold for *c*

Given the model fit, the overall false positive rate for the mutations detected was calculated as follows:

1. Compute FPR(c) = E(c)/[E(c) + T(c)]. This gives the false positive rate at each composite coverage value.
2. Compute the number of false positive mutations in the scsSNV set as a function of *c* as **F**_c_ = [FPR(c)] **M**_c_
3. Compute the expected aggregate false positive rate when thresholding at *c*\* as:

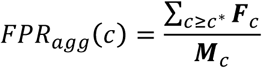
4. Choose *c* such that the aggregate false positive rate is controlled under some consensus value (e.g. 10%)

## References

1. Islam, S. et al. Quantitative single-cell RNA-seq with unique molecular identifiers. Nat Meth 11,163–166 (2013).

2. Leung, M. L., Wang, Y., Waters, J. & Navin, N. E. SNES: single nucleus exome sequencing. Genome Biology 16,55–10 (2015).

3. Xu, X. et al. Single-Cell Exome Sequencing Reveals Single-Nucleotide Mutation Characteristics of a Kidney Tumor. Cell 148,886–895 (2012).

4. Hou, Y. et al. Single-Cell Exome Sequencing and Monoclonal Evolution of a JAK2 Negative Myeloproliferative Neoplasm. Cell 148,873–885 (2012).

5. Baslan, T. et al. Genome-wide copy number analysis of single cells. Nat Protoc 7,1024–1041 (2012).

6. Lodato, M. A. et al. Somatic mutation in single human neurons tracks developmental and transcriptional history. Science 350,94–98 (2015).

7. Zong, C., Lu, S., Chapman, A. R. & Xie, X. S. Genome-Wide Detection of Single-Nucleotide and Copy-Number Variations of a Single Human Cell. Science 338,1622–1626 (2012).

8. Dean, F. B. et al. Comprehensive human genome amplification using multiple displacement amplification. Proceedings of the National Academy of Sciences 99,5261–5266 (2002).

9. Huang, L., Ma, F., Chapman, A., Lu, S. & Xie, X. S. Single-Cell Whole-Genome Amplification and Sequencing: Methodology and Applications. Annu. Rev. Genom. Hum. Genet. 16,79–102 (2015).

10. Gawad, C., Koh, W. & Quake, S. R. Single-cell genome sequencing: current state of the science. Nature Publishing Group 17,175–188 (2016).

11. Esteban, J. A., Salas, M. & Blanco, L. Fidelity of phi 29 DNA polymerase. Comparison between protein-primed initiation and DNA polymerization. J. Biol. Chem. 268,2719–2726 (1993).

12. Fryxell, K. J. & Zuckerkandl, E. Cytosine deamination plays a primary role in the evolution of mammalian isochores. Molecular Biology and Evolution 17,1371–1383 (2000).

13. Lindahl, T. & Nyberg, B. Heat-induced deamination of cytosine residues in deoxyribonucleic acid. Biochemistry 13,3405–3410 (1974).

14. Frederico, L. A., Kunkel, T. A. & Shaw, B. R. A sensitive genetic assay for the detection of cytosine deamination: determination of rate constants and the activation energy. Biochemistry 29,2532–2537 (1990).

15. McKenna, A. et al. The Genome Analysis Toolkit: a MapReduce framework for analyzing next-generation DNA sequencing data. Genome Res. 20,1297–1303 (2010).

16. Hoang, M. L. et al. Genome-wide quantification of rare somatic mutations in normal human tissues using massively parallel sequencing. Proceedings of the National Academy of Sciences 113,9846–9851 (2016).

17. Cibulskis, K. et al. Sensitive detection of somatic point mutations in impure and heterogeneous cancer samples. Nature Biotechnology 31,213–219 (2013).

18. Koboldt, D. C. et al. VarScan: variant detection in massively parallel sequencing of individual and pooled samples. Bioinformatics 25,2283–2285 (2009).

19. Supek, F. & Ben Lehner. Clustered Mutation Signatures Reveal that Error-Prone DNA Repair Targets Mutations to Active Genes. Cell 170, 534.e1–534.e23 (2017).

20. Crouse, G. F. Non-canonical actions of mismatch repair. DNA Repair 38,102–109 (2016).

21. Zafar, H., Wang, Y., Nakhleh, L., Navin, N. & Chen, K. Monovar: single-nucleotide variant detection in single cells. Nat Meth 13,505–507 (2016).

22. Roth, A. et al. Clonal genotype and population structure inference from single-cell tumor sequencing. Nat Meth 13,573–576 (2016).

## References

1. McKenna, A. et al. The Genome Analysis Toolkit: a MapReduce framework for analyzing next-generation DNA sequencing data. Genome Res. 20,1297–1303 (2010).

2. Auton, A. et al. A global reference for human genetic variation. Nature 526,68–74 (2015).

3. Marchini, J. et al. Integrating sequence and array data to create an improved 1000 Genomes Project haplotype reference panel. Nature Communications 5, 3934 (2014).

